# A Framework for Automated Construction of Heterogeneous Large-Scale Biomedical Knowledge Graphs

**DOI:** 10.1101/2020.04.30.071407

**Authors:** Tiffany J. Callahan, Ignacio J. Tripodi, Lawrence E. Hunter, William A. Baumgartner

**Affiliations:** Computational Bioscience Program, University of Colorado Anschutz Medical Campus, CO, USA; Computer Science Department, University of Colorado Boulder, CO, USA

## Abstract

**Motivation:** Although knowledge graphs (KGs) are used extensively in biomedical research to model complex phenomena, many KG construction methods remain largely unable to account for the use of different standardized terminologies or vocabularies, are often difficult to use, and perform poorly as the size of the KG increases in scale. We introduce PheKnowLator (Phenotype Knowledge Translator), a novel KG framework and fully automated Python 3 library explicitly designed for optimized construction of semantically-rich, large-scale biomedical KGs. To demonstrate the functionality of the framework, we built and evaluated eight different parameterizations of a large semantic KG of human disease mechanisms. PheKnowLator is available at: https://github.com/callahantiff/PheKnowLator.

## 1 INTRODUCTION

Knowledge graphs (KGs) facilitate the representation of relationships among heterogeneous data types and have been used extensively in biomedical research to model complex biological phenomena [1]. In the biomedical domain, KGs are usually constructed from a combination of ontologies, linked open data, and experimental data [2–5]. Methods to construct KGs vary from domain-specific manual approaches [6] to semi-supervised rule-based [7] and fully automated unsupervised approaches [8]. To date, the vast majority of KG construction algorithms have been developed in order to create more manageable representations of large free-text corpora (e.g. scientific articles) [9,10], to derive novel associations between existing concepts [11,12], and add evidence to existing systems or KGs [13,14]. While many data-driven KG construction methods have been developed, they remain largely unable to automatically construct KGs from multiple disparate data sources, combine KGs created by different systems, and collaborate or share KGs across institutions due to their inability to account for the use of different schemas, standards, and vocabularies [15].

Used extensively in life sciences research, the Semantic Web was created to resolve these types of knowledge integration problems [16]. The Web Ontology Language (OWL) is a Semantic Web standard for graph-based knowledge representation and reasoning [17]. OWL is highly expressive, enabling the integration of heterogeneous data using explicit semantics, and allows for the generation of new knowledge using deductive logic. Unfortunately, existing OWL-based KG construction methods are often built using complicated programs or toolsets, are written in nonstandard programming languages, and can require extensive ontological expertise and computational resources.

In addition to technical challenges, there are multiple approaches to modeling knowledge in graphs, each with their own advantages (and adherents). For example, there are alternative approaches to the quantification implied by an edge between two nodes or in restrictions on the introduction of cycles in a graph. Rather than choosing one approach, we propose a more robust solution in the form of a generalizable tool equipped with functionality to accommodate a wide range of use cases. To accomplish this goal, while solving some of the aforementioned technological challenges, we introduce PheKnowLator (Phenotype Knowledge Translator), a fully automated Python 3 library explicitly designed for optimized construction of semantically-rich, large-scale biomedical KGs.

## 2 METHODS

### 2.1 PheKnowLator Framework

PheKnowLator requires three user-customizable inputs to construct a KG: a list of ontologies, a list of linked open data and/or non-ontology data sources, and source-specific edge specifications. To reduce user burden, PheKnowLator includes a universal file parser to facilitate the integration of new data sources and suggests default input parameters. Utilizing these inputs, a KG is constructed via three steps: (1) ontology and data source download; (2) preprocess data (i.e. filter data, map identifiers, and merge ontologies) and construct edge lists; and (3) construct KGs and property graphs, and output KG metadata.

#### 2.1.1 Build Types

PheKnowlator was designed to run from two different development stages: (1)<u>Full</u>: for users who want to build a KG from scratch and (2) <u>Post-Closure</u>: for users who want to apply PheKnowLator functionality to a pre-existing KG (e.g. KGs that have been reasoned or used in other learning tasks).

#### 2.1.2 Construction Approaches

PheKnowLator is equipped with the two most common KG representation approaches utilized by the Semantic Web community: instance-based [4] and subclass-based [3]. For the instance-based method, non-ontology data (e.g. Reactome and the Comparative Toxicogenomic Database) are connected to an instance of an existing ontology class. For the subclass-based method, non-ontology data are connected by subclassing an existing ontology class. Both methods require a map between non-ontology entities and ontology classes.

#### 2.1.3 Relations

It is assumed that relations will be drawn from ontologies (e.g. the Relations Ontology). By leveraging ontology structure, PheKnowLator is able to offer users the option of building a KG with or without inverse relations. Additionally, it can automatically generate inverse relations for one-way, implicitly symmetric relations for edge types that represent different kinds of biological interactions (e.g. *molecularly interacts with*).

#### 2.1.4 Creation of Property Graphs

OWL is a highly expressive representation language, but its use comes at the cost of a structurally complex KG. Many of the triples responsible for this complexity do not contain biologically meaningful information (i.e. OWL-encoded triples). To create a version of a PheKnowLator KG that only includes biologically meaningful triples (i.e. all OWL-encoded triples have been removed), we extend OWL-NETS [18], a statistical learning method which losslessly removes triples containing OWL semantics by automatically decoding KGs in two independent steps: (1) removing all triples that connect non-biologically meaningful subject and object nodes; and (2) decoding owl-encoded classes built using *owl:intersectionOf, owl:unionOf*, and *owl:Restriction* constructs. For more details, please see the Wiki^1^.

### 2.2 Technical Specifications

PheKnowLator was developed using Python 3.6.2 and is available through PyPI. The library is dependent on OWLTools v0.3.0^2^. Additionally, we provide several out-of-the-box tools for building a KG, including a Docker image and a Jupyter Notebook. Docker improves reproducibility by containerizing the specific version of software and data used for each KG build.

### 2.3 Preliminary Evaluation

A biologically meaningful KG, i.e. one where the relations between nodes accurately reflect the intent of the data stored in the original sources, was developed by working with a PhD-level molecular biologist (Figure 1).

**Figure 1.**
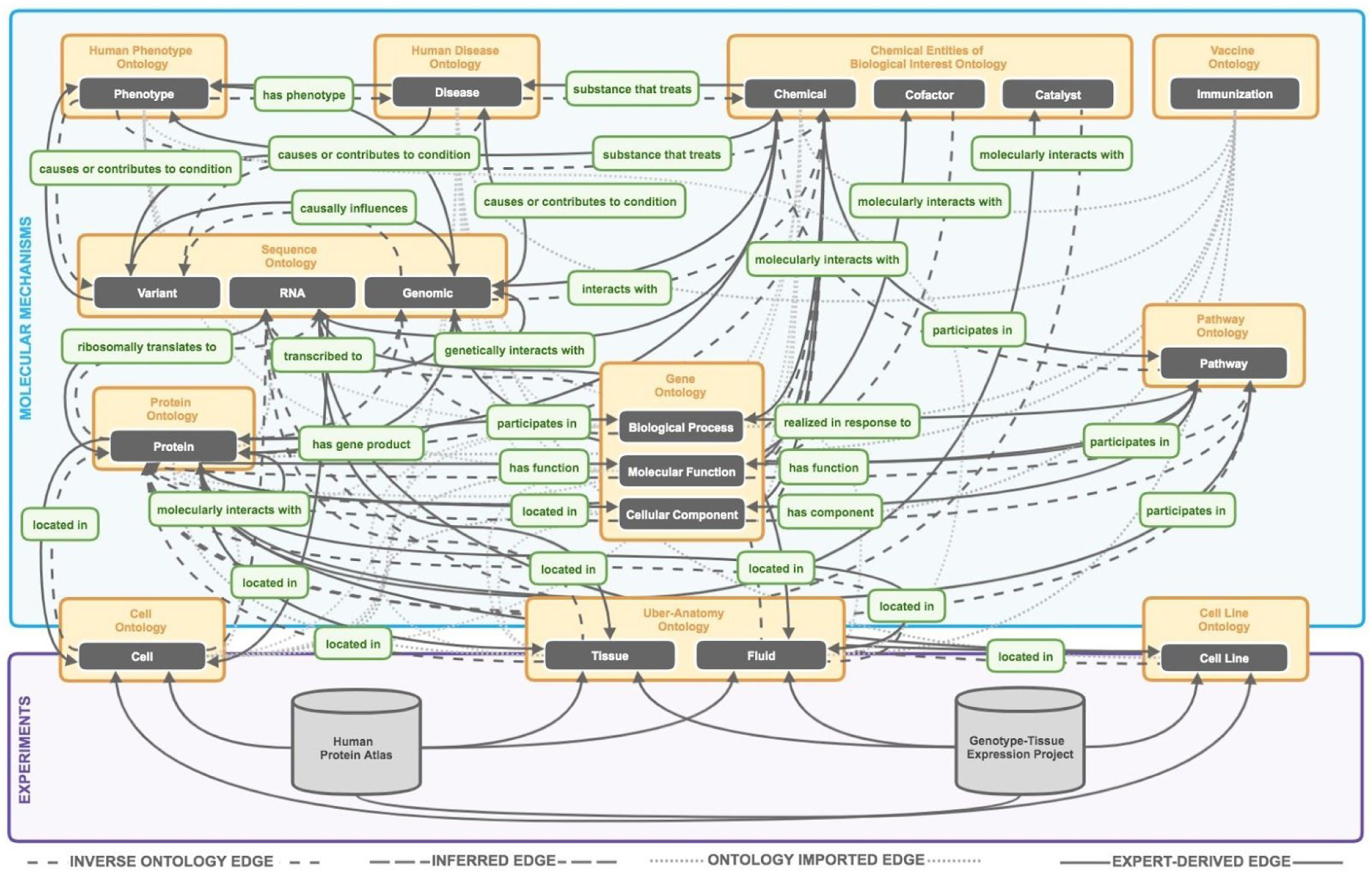
The knowledge representation used to construct the eight different parameterizations of the PheKnowLator KGs. The purple box represents experimental data and the blue box represents molecular mechanisms created by integrating open biomedical ontologies and linked open data. Experimental data was taken from the Human Protein Atlas and the Genotype-Tissue Expression Project (GTEx). Molecular mechanisms were created from 12 OBOs, which when joined with the 24 linked open data sources, facilitated the following 34 distinct edge types: chemical-disease, chemical-gene, chemical-biological process, chemical-cellular component, chemical-molecular function, chemical-pathway, chemical-phenotype, chemical-protein, chemical-rna, disease-phenotype, gene-disease, gene-gene, gene-pathway, gene-phenotype, gene-protein, gene-rna, biological process-pathway, pathway-cellular component, pathway-molecular function, protein-anatomy, protein-catalyst, protein-cell, protein-cofactor, protein-biological process, protein-cellular component, protein-molecular function, protein-pathway, protein-protein, rna-anatomy, rna-cell, rna-protein, variant-disease, variant-gene, and variant-phenotype. Finally, the green boxes represent relations from the Relations Ontology. For additional information on the data (including preprocessing and filtering) used to construct the KG, please see the project Wiki (https://github.com/callahantiff/PheKnowLator/wiki/v2-Data-Sources).

#### 2.3.1 Data Sources

The KGs were constructed using 12 open biomedical ontologies, twenty-four linked open datasets, and results from two large-scale, experimentally-derived datasets. Additional information on these resources is provided on the Wiki^1^ (see V2.0.0).

#### 2.3.3 Parameterizing KGs

Eight different KGs were constructed in order to demonstrate the range of KGs that PheKnowLator can produce. These KGs were built by varying the construction approach (i.e. instance- vs. subclass-based), relations (i.e. using only relations vs. relations with their inverse), and whether or not OWL-encoded classes were decoded. All KGs were constructed on a CentOS Linux server 7.0.1406 equipped with 24 CPUs (X5680 @ 3.33GHz) and 98 GB of memory.

#### 2.3.3 Metrics

Metrics to evaluate each of the eight PheKnowLator KG parameterizations included: (1) <u>Performance</u>: run-time (minutes), memory use (GB), and average load factor at one minute were recorded for each of the KG build steps described in Section 2.1. For Steps 1-2, which are the same across all KG parameterizations, the minimum, average, and maximum runtime were calculated. Additionally, Step 3 was run once per KG parameterization. For all steps, average load factor at one minute and memory use were calculated from system statistics which were output every 60 seconds during runtime; (2) <u>D</u> <u>escriptives</u>: counts of classes, axioms, and individuals were obtained using the *info* command from OWLTOOLs.

## 3 RESULTS

On average, Step 1 (i.e. downloading ontology and linked open data) took 9.08 minutes (7.31-11.07 minutes), used an average of 0.57 GB of memory (0.20-1.26 GBs), and had an average load factor at one minute of 6.09 processes (4.79-6.74 processes). Step 2 (i.e. preprocessing data and constructing edge lists) took an average of 21.27 minutes to complete (16.29-23.70 minutes), used an average of 1.32 GB of memory (0.05-2.95 GBs), and had an average load factor at one minute of 3.01 processes (2.05-7.51 processes). The performance metrics for Step 3 (i.e. constructing KGs) for each KG parameterization are shown in Figure 2. In general, these plots suggest that on average, KG parameterizations constructed using the subclass construction approach take longer to run, use more memory to build, and have a higher average factor load at one minute than KGs built using the instance construction approach. Regardless of the construction approach, KGs that included inverse relations and performed OWL decoding took longer to run, used more memory, and had a higher average load factor at one minute.

**Figure 2.**
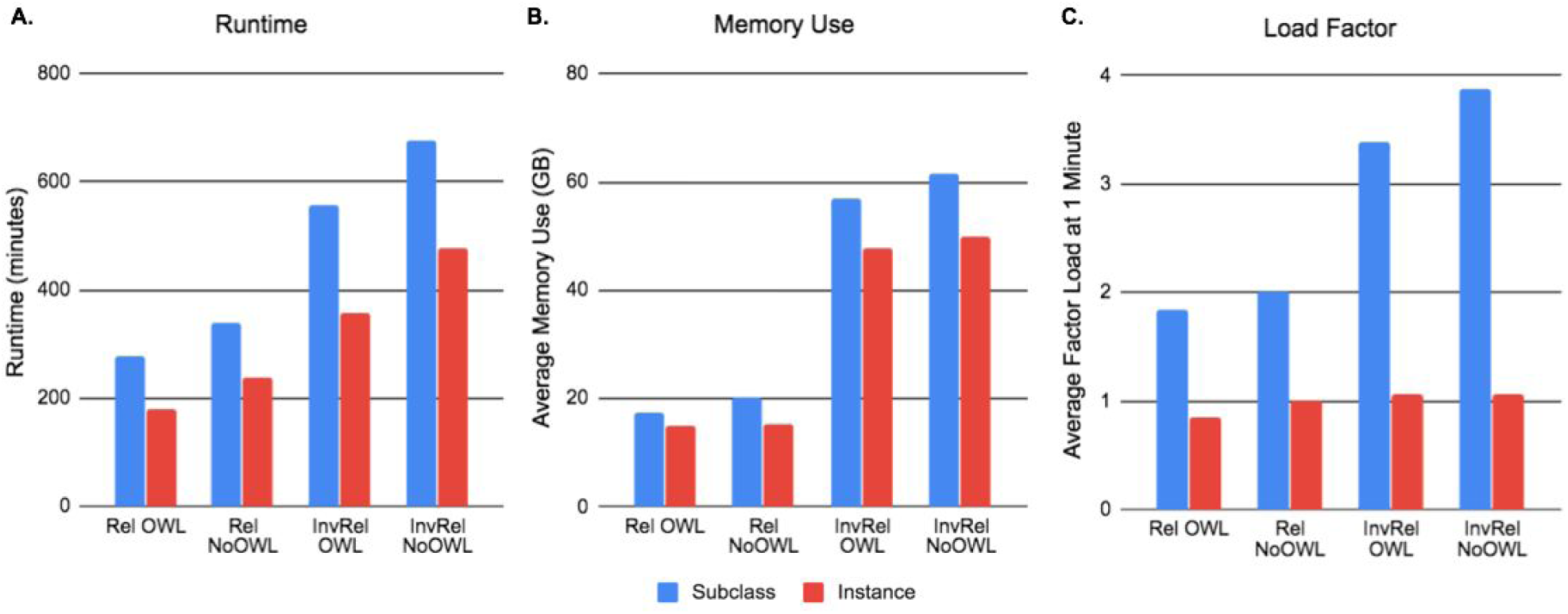
KG parameterization performance metrics (a: runtime, b: average memory use, and c: average load factor at one minute) for the KG construction step of the PheKnowLator algorithm. In all plots, blue bars represent the subclass construction approach and red bars represent the instance construction approach. The x-axis features different combinations of parameterizations of the relations (i.e. Rel: relations only vs InvRel: relations and inverse relations) with or without decoding OWL semantics.

Descriptives of each KG parameterization are shown in Table 1. All KG parameterizations were built on the same base set of merged ontologies (downloaded in Step 1 and merged in Step 2; Figure 1) which contained 366,847 classes, 3,923,630 axioms, 825 object properties, and 123 individuals. As demonstrated by Table 1, KGs built using the subclass construction approach tended to be larger (i.e. contained more classes, resulted in larger file size, and contained more triples) than instance-constructed KGs. This supports the longer runtimes and use of more resources for the subclass-constructed-KGs, observed above.

**Table 1.** Descriptives for each PheKnowLator KG parameterizations.

| Construction Approach | Relations | Decoding OWL | Axioms | Classes | Individuals |
| --- | --- | --- | --- | --- | --- |
| Subclass | Relations Only | Yes | 11,635,203 | 0 | 0 |
|  |  | No | 16,570,590 | 966,961 | 123 |
|  | Relations + Inverse | Yes | 20,442,012 | 0 | 0 |
|  |  | No | 24,418,347 | 966,961 | 123 |
| Instance | Relations Only | Yes | 13,801,188 | 0 | 0 |
|  |  | No | 16,815,593 | 366,847 | 667,826 |
|  | Relations + Inverse | Yes | 22,497,531 | 0 | 0 |
|  |  | No | 24,552,884 | 366,847 | 667,826 |

Relations only: single directed edge; Relations + Inverse: relations and inverse relations via applying *owl:inverseOf* for each included relation.

## 4 DISCUSSION

We introduce PheKnowlator, the first fully customizable KG construction framework enabling users to build complex KGs that are Semantic Web compliant and amenable to automatic OWL reasoning, conform to contemporary property graph standards, and are importable by today’s popular graph toolkits. PheKnowlator provides this functionality by offering multiple build types, can automatically include inverse edges, creates OWL-decoded KGs to support automated deductive reasoning, and outputs KGs in several formats (e.g. triple edge lists, OWL API-formatted RDF/XML and graph-pickled MultiDiGraph). By providing flexibility in the way relations are modeled and facilitates the creation of property graphs, PheKnowLator enables the use of cutting edge graph-based learning and sophisticated network inference algorithms.

Our preliminary results evaluated different parameterizations of PheKnowLator KGs on a variety of dimensions. PheKnowLator KGs have also recently been used in applications of toxicogenomic mechanistic inference [14] and biomedical hypergraphs [19] and we’d like to invite the community to collaborate with us to examine the utility of PheKnowLator across a wider variety of use cases. Although not included in this preliminary evaluation, we are in the process of running different reasoners [20] over each KG parameterization. Once complete, the accuracy of the newly inferred triples, for each KG, will be reviewed by both clinical and biological domain experts. Finally, we are in the process of setting up a cloud-based Neo4j instance which will be made publicly available.

## ACKNOWLEDGEMENTS

This work was supported by a Training Grant from the NLM, NIH to the University of Colorado Biomedical Informatics Training Program T15LM009451. We’d like to thank Adrianne Stefanski for consulting on the KG knowledge representation, David Farrell and Luca Cappelletti for their technological expertise and assistance with continuous integration and Docker, and Jordan Wrywa for content review.

## Footnotes

1 https://github.com/callahantiff/PheKnowLator/wiki

2 https://github.com/owlcollab/owltools

## Notes

### Competing Interest Statement

The authors have declared no competing interest.

